# Bridging energy and ribosomal allocation models to predict the cost of traits in different environments

**DOI:** 10.1101/2025.09.30.679561

**Authors:** Mohammadjavad Meghrazi, Sarah P. Otto

## Abstract

Many traits are costly because they require the diversion of resources from cell reproduction, however, the effect of environmental conditions and genetic background on the cost of traits is not well understood. Two different frameworks have been proposed to quantify the resource costs of traits, focusing on either energy allocation or ribosome allocation. These frameworks implicitly assume energy provisioning or protein production limits growth, respectively, but the connection between the two limitations has been underexplored. To better connect these frameworks, we reformulate previous models and incorporate the degradation, recycling, and energetic demands for cell maintenance to quantify the cost of traits, depending on the nature of the resources diverted, genetic background, and the environmental conditions experienced. Notably, our model predicts that increasing food quality increases the cost of traits that require the production of new structures, while decreasing the cost of traits requiring energy expenditure. Understanding how environmental change affects the cost of traits has important implications for the evolution of various traits, including antimicrobial resistance. Moreover, the model also accounts for several aspects of the observed relationship between the macromolecular composition of the cells and growth rate (also known as bacterial growth laws).

**Author Summary:** Many traits divert resources from reproduction and are costly due to the physiological trade-offs organisms face. These traits might require energy expenditure or production of macromolecules, and it is unclear how their costs can be compared given their different units. Moreover, the effect of environmental conditions on the cost of traits is not well understood. Here, we develop a framework that considers limitations in energy provisioning and protein production simultaneously and enables us to compare the cost of traits requiring different resources. Using this model, we explore how environmental quality and genetic background affect the cost of traits.

## Introduction

Traits underlying adaptation to a new environment sometimes require diverting resources from reproduction towards other functions: neutralizing toxins, moving macromolecules, buffering environmental changes, etc. Due to physiological trade-offs, the diversion of resources can reduce the reproduction rate, and this reduction can be defined as the resource cost of traits (hereafter cost of traits)[9]. Different traits may require the allocation of different resources. For example, resistance to antibiotics could be achieved by the production of new proteins (e.g., proteins that degrade the antibiotics) or energy expenditure (e.g., activating pumps that remove antibiotics from the cytoplasm)[17]. Both of these traits are costly, but it is unclear how their costs, i.e., protein production and energy expenditure, can be compared given their different units. Moreover, the cost depends on the environment and genetic background of the cells in a manner that is not well understood[11]. As less costly traits are more likely to evolve all else being equal, if we can convert the costs of different traits to a common unit and compare them, we can predict which traits are more likely to evolve. Moreover, understanding how the environment and genetic background affect the cost of traits will help us predict the conditions that facilitate the evolution of various traits, including resistance to stressors (e.g., antibiotics, extreme temperatures, and predation). This can have important implications for the management of the antimicrobial resistance crisis and, more broadly, understanding the evolutionary response of organisms to the increased prevalence of human-induced stressors.

Energy allocation and ribosome allocation are the two main frameworks used to study the cost of traits, however, they have different underlying assumptions, and the connection between them is underexplored[16, 24, 21, 4, 2]. According to energy allocation models, the cost of a trait is translated into energy requirements and depends on the fraction of the cell’s energy budget consumed[16]. This is equivalent to assuming cells are energy-limited, i.e., they cannot increase their energy budget[19]. On the other hand, the ribosome allocation framework considers protein production to be the limiting process in cell reproduction, with emphasis on limitation in translation. Therefore, allocating proteins to a trait decreases the allocation to the protein production machinery, which increases the time required to reproduce the proteome of a cell[28, 23, 14]. Energy provisioning and translation are specific cases of processing nutrients (provisioning) and converting them to macromolecules (biosynthesis), respectively (Figure 2). As there is ample evidence for limitations in both provisioning and biosynthesis, the unification of these two frameworks is necessary for a better understanding of the constraints under which cells grow and reproduce[13].

We develop a macromolecule allocation framework that considers provisioning and biosynthesis simultaneously to quantify how each limits cell growth. Using this model, we quantify the cost of allocating different resources to traits and the effect of environmental conditions and genetic background on this cost. Notably, we show that the cost of traits that require macromolecule production is higher in environments favorable for growth. However, the cost of traits that require energy expenditure or a higher degradation rate of harmful proteins depends on the extent to which provisioning and biosynthesis limit growth, which in turn depends on genotype and environment. Factors that increase the speed of provisioning and make it less limiting (e.g., higher food quality) decrease the cost of energy expenditure and increase the cost of degradation, while factors that increase the speed of biosynthesis (e.g., faster ribosomes) have the opposite effect. Moreover, our model shows how degradation, recycling, and energetic demands for maintenance simultaneously affect the relationship between the fraction of ribosomes in the proteome and growth rate (also known as bacterial growth laws; explained in Box 1), particularly during slow growth.

#### Box1: Bacterial growth laws

The macromolecular composition of single-celled organisms is tightly correlated with their growth rate at steady state[22]. When food quality is altered, a positive linear relationship is observed between the growth rate and the fraction of ribosomes in the proteome, which is often measured approximately using the ratio of RNA to proteins (Figure 1a)[23]. However, when ribosomes are partially inhibited, the relationship between growth rate and the ribosomal fraction switches to a negative one[23]. Both of these relationships between the fraction of ribosomes and growth rate are robust and linear, except at slow growth, where the ribosomal fraction is higher than expected based on the linear relationship (concave up)[7]. These patterns are called bacterial growth laws[23] (Figure 1a). More recently, these trends have been confirmed by directly measuring the fraction of ribosomes in the proteome [18, 12].

Scott et al. (2010)[23] used a three-compartment model of the proteome to account for bacterial growth laws. In this model, ribosomes and other proteins involved in translation are called the R fraction (*ϕ*_*R*_), metabolic enzymes that provide energy and substrates for ribosomes are called the P fraction (*ϕ*_*P*_), and the remaining proteins (mainly structural and maintenance proteins), which are assumed to have a fixed relative abundance, are called the fixed fraction *ϕ*_*Q*_. The cell can adjust the ratio of R and P fractions to maximize protein production rate, which they assumed is tightly linked to growth rate. Therefore when the cell is placed in low-quality food, which requires more enzymes to be processed, the P fraction is expected to increase. In the presence of ribosome inhibitors, however, the R fraction must rise to compensate for the inhibition (Figure 1d).

From the perspective of Scott et al. (2010), the growth rate equals the ribosomal fraction times the protein production rate per ribosome mass (i.e., the average speed of ribosomes). To explain the linear relationship between the ribosomal fraction and growth rate when altering food quality, ribosomes are assumed to work at a constant speed, regardless of the conditions within the cell[23]. Otherwise, if the speed of ribosomes covaries with the ribosomal fraction, the relationship between the ribosomal fraction and growth rate would not be linear (Figure 1b, blue line). Nevertheless, it is known that the speed of active ribosomes is positively correlated with food quality and approaches half of its maximal value as the growth rate approaches zero (Figure 1c)[8, 27, 20, 7]. Moreover, if the speed of ribosomes were always constant, then the ribosomal fraction is expected to decline to zero as the growth rate approaches zero, which is not observed (Figure 1b, red line)[18].

To explain these discrepancies, previous studies have investigated the role of inactive ribosomes, degradation, and variable speed of ribosomes [7, 18, 5, 15, 6]. To our knowledge, none of the previous studies have investigated the effects of the recycling of degraded macromolecules and energetic demands for maintenance (hereafter maintenance demands), which are particularly important during slow growth[16].

**Figure 1.**
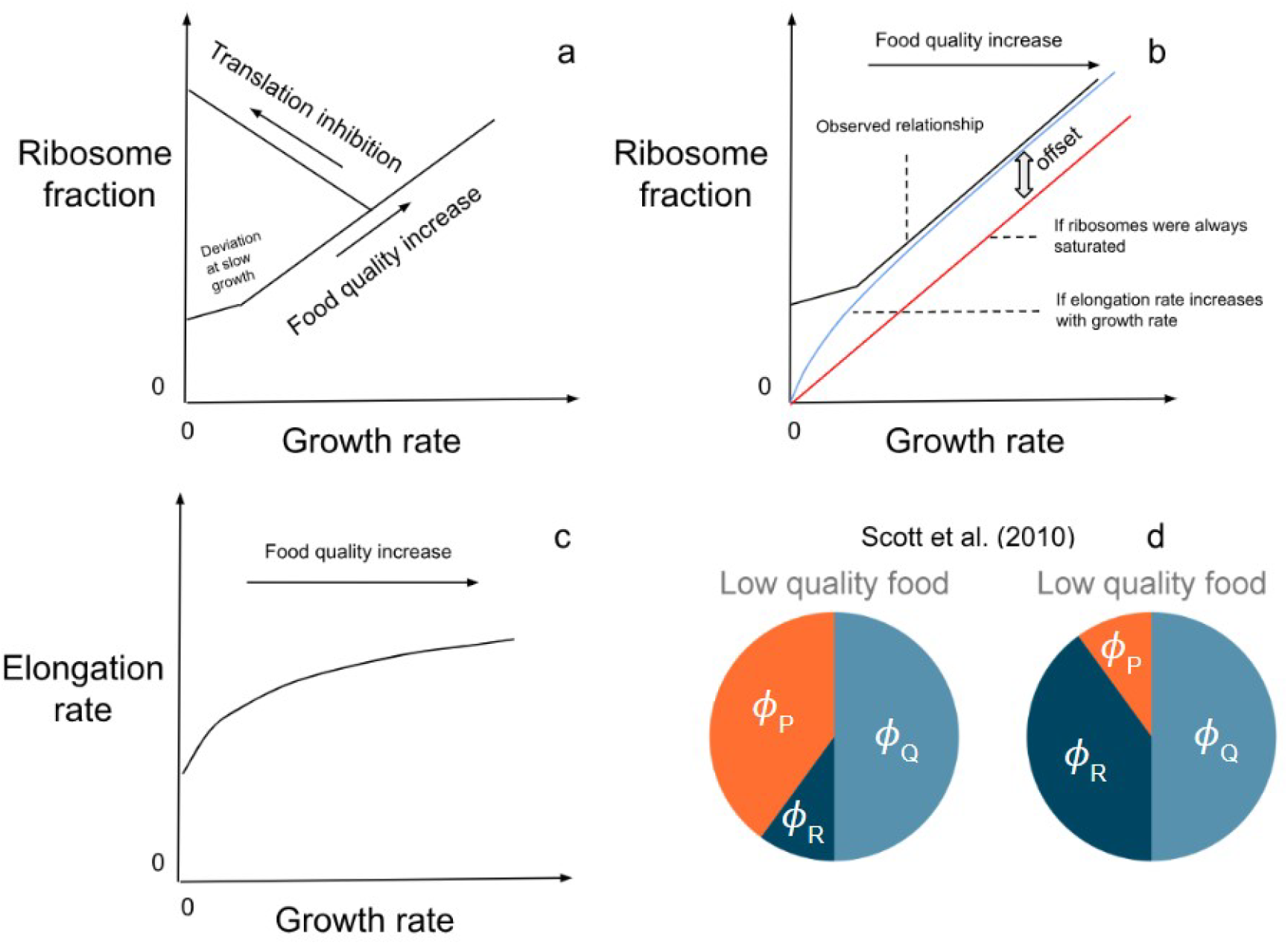
Bacterial growth laws describe the relationship between growth rate and the fraction of ribosomes in the proteome when altering food quality or inhibiting translation (panel a). When altering food quality, this relationship has a positive y-intercept (panel b, black line), which is not expected if ribosomes were always working at a constant speed (red line)[18]. On the other hand, if the speed of ribosomes increases with growth rate, a decelerating relationship between growth rate and ribosomal fraction is expected (blue line), which is not observed[6]. Nevertheless, the speed of ribosomes is positively correlated with food quality and approaches half of its maximum value as the growth rate of cells approaches zero (panel c)[7]. To explain the bacterial growth laws, Scott et al. (2010) used a three-compartment model of the proteome (panel d). In their model, when food quality is low, the cell increases allocation to the metabolic proteins that supply ribosomes (*ϕ*_*P*_). On the other hand, when ribosomes are inhibited, the cell increases the ribosomal fraction (*ϕ*_*R*_) to compensate for the inhibition. The fixed fraction (*ϕ*_*Q*_) is composed of proteins with constant relative abundance.

**Figure 2.**
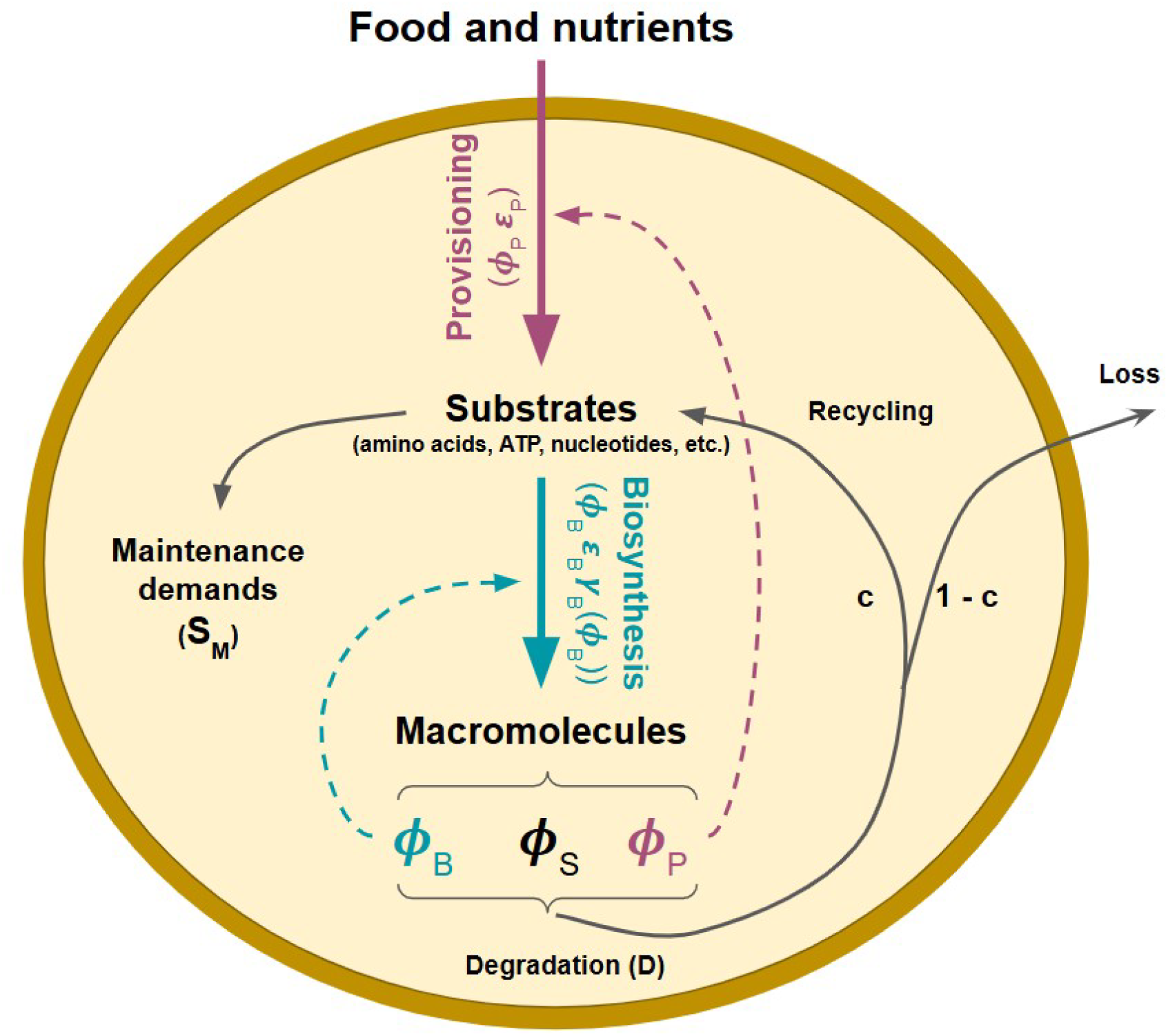
We divide macromolecules into three fractions: The provisioning fraction (*ϕ*_*P*_), the biosynthesis fraction (*ϕ*_*B*_), and the structural fraction (*ϕ*_*S*_). Specifically, the provisioning fraction (*ϕ*_*P*_) converts food and nutrients into substrates (i.e., energy stores like ATP, amino acids, nucleotides, etc.) and consists of enzymes that break down food, produce various substrates, and the uptake pumps and channels. Substrates are then used for both maintenance tasks (*S*_*M*_) and macromolecule production by the biosynthesis fraction (*ϕ*_*B*_), which consists of ribosomes, DNA and RNA polymerases, membrane and cell wall synthesis enzymes, and other macromolecules involved in polymerization. In addition, macromolecules tightly correlated in abundance with ribosomes or other biosynthesis enzymes, e.g., tRNAs, and mRNAs, are part of this fraction [15, 3]. The structural fraction (*ϕ*_*S*_) involves all the other macromolecules not directly involved in facilitating provisioning or biosynthesis, including DNA, membrane lipids, cell wall, storage macromolecules, half-made macromolecules, and proteins that help with structure, degradation, and maintenance. Degradation (*D*) breaks down macromolecules and partially recycles constituent substrates. Solid lines represent biomass flux, and dashed lines represent the enzymatic facilitation of biomass flux. The mass-standardized flux (flux in a cell divided by the mass of cell macromolecules) of different pathways is written inside parentheses.

### Model

To quantify the rate of reproduction (i.e., biomass production), we develop a three-compartment model similar to Scott et al. (2010) (described in box 1) but with three differences[23]. First, we include all macromolecules in our model rather than just proteins. Second, the speed of biosynthesis machinery is allowed to vary. Third, we incorporate degradation, recycling, and energetic demands for maintenance.

We divide cell macromolecules into three fractions based on their metabolic role. The macromolecules that uptake food and nutrients and convert them into substrates (i.e., energy stored in different forms, amino acids, nucleotides, etc.) make up the provisioning fraction (*ϕ*_*P*_). The macromolecules that synthesize new macromolecules using substrates comprise the biosynthesis fraction (*ϕ*_*B*_). The remaining macromolecules are grouped into the structural fraction (*ϕ*_*S*_) (Figure 2). See more details on the components of each fraction in S1 Text. We assume that the structural fraction is fixed but the cell can optimize the ratio allocated to provisioning versus biosynthesis to maximize biomass production.

Three processes affect the flux of substrates supplied to the substrate pool. 1-The provisioning fraction supplies substrates proportional to its mass at the rate *M*_*P*_ *ϵ*_*P*_, where *M*_*P*_ is the mass of the provisioning machinery and *ϵ*_*P*_ (provisioning efficiency) is the substrate supply rate per unit mass of the provisioning machinery. 2-Maintenance demands consume substrates (mainly in the form of energy) at a constant rate equal to *M*_*Total*_ *S*_*M*_, where *S*_*M*_ is the substrate demand for maintenance per unit mass of cell macromolecules and *M*_*Total*_ is the total mass of cell macromolecules. 3-Recycling of degraded macromolecules supplies substrates proportional to the degradation rate and the efficiency of substrate recycling at the rate *M*_*Total*_ *c D*, where *D* is the degradation rate per unit of macromolecule mass and *c* (recycling efficiency) is the fraction of substrates recycled after degradation. Overall, substrates are supplied to the substrate pool at the rate *M*_*Total*_(*ϕ*_*P*_ *ϵ*_*P*_ −*S*_*M*_ + *c D*), where *ϕ*_*P*_ = *M*_*P*_ */M*_*Total*_. The biosynthesis fraction consumes substrates proportional to the mass and speed of biosynthesis machinery at the rate *M*_*B*_ *ϵ*_*B*_ *γ*_*B*_(*ϕ*_*B*_), where *M*_*B*_ is the mass of biosynthesis machinery, *ϵ*_*B*_ (biosynthesis efficiency) is the rate of substrate consumption per unit mass of biosynthetic machinery when saturated, and *γ*_*B*_(*ϕ*_*B*_) is the biosynthesis saturation level, which depends on the steady-state substrate concentration. As the steady-state substrate concentration depends on the biosynthesis fraction, we model the biosynthesis saturation level directly as a function of the biosynthesis fraction rather than the steady-state substrate concentration. Moreover, substrates are allocated to new cells in proportion to the birth rate and the steady-state substrate concentration (dilution due to growth, see S1 Text for more details). Table 1 summarizes the notation.

**Table 1:** List of variables and their units.

| Variable symbol | Description | Unit |
| --- | --- | --- |
| $M_X$ | Mass of macromolecules in fraction $X$ ($X = P, B$, or $S$) | g |
| $M_{Total}$ | Total mass of cell macromolecules | g |
| $\phi_X$ | The size of fraction $X$, which equals $\frac{M_X}{M_{Total}}$<br>($X = P, B$, or $S$) | dimensionless |
| $\epsilon_P$ | Provisioning efficiency: mass of substrates produced by provisioning machinery, per unit mass of provisioning machinery, per unit time | $h^{-1}$ |
| $\epsilon_B$ | Biosynthesis efficiency: mass of substrates consumed (or macromolecules produced) by biosynthetic machinery when saturated, per unit mass of biosynthetic machinery, per unit time | $h^{-1}$ |
| $\gamma_B(\phi_B)$ | The saturation level of biosynthesis which is a function of biosynthesis fraction. | dimensionless |
| $b$ | Birth rate: the mass of macromolecules produced per unit mass of macromolecules, per unit time | $h^{-1}$ |
| $D$ | Degradation rate: mass of macromolecules degraded, per unit mass of macromolecules, per unit time | $h^{-1}$ |
| $S_M$ | Maintenance demands: mass of substrate consumed, per unit mass of macromolecules, per unit time | $h^{-1}$ |
| $\alpha$ | The maximum substrate supply rate (when the provisioning fraction is maximal), which equals $\epsilon_P(1 - \phi_S) - S_M + c D$ | $h^{-1}$ |
| $A$ | The extent to which provisioning is limiting, which equals $\frac{\epsilon_B}{(\sqrt{\epsilon_B} + \sqrt{\epsilon_P})^2}$. When provisioning is the limiting step, $A$ approaches 1, and when biosynthesis is limiting, $A$ approaches 0. | dimensionless |
| $R$ | The return on investment, which equals $\frac{\epsilon_P \epsilon_B}{(\sqrt{\epsilon_B} + \sqrt{\epsilon_P})^2}$. $R$ equals the birth rate achieved in the absence of degradation and maintenance demands if all macromolecules were allocated to provisioning and biosynthesis ($\phi_S = 0$).<br>Notice that $\frac{1}{\sqrt{R}} = \frac{1}{\sqrt{\epsilon_B}} + \frac{1}{\sqrt{\epsilon_P}}$. | $h^{-1}$ |

We assume the macromolecule production rate determines the rate of cell reproduc-tion. Therefore the birth rate (*b*) equals the net macromolecule production rate per unit of macromolecule mass, which equals the biosynthesis rate minus macromolecule degradation rate divided by the mass of all macromolecules. Assuming biosynthesis rate equals the rate of substrate consumption by the biosynthesis fraction (i.e., no mass lost during biosynthesis), the birth rate equals:

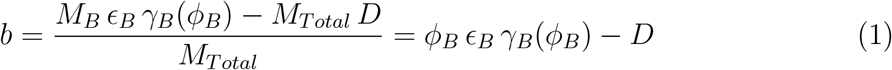

Assuming the degradation rate is independent of macromolecular allocation, the allocation that maximizes *ϕ*_*B*_ *ϵ*_*B*_ *γ*_*B*_(*ϕ*_*B*_) will maximize the birth rate. However, there is a trade-off between the biosynthesis fraction (*ϕ*_*B*_) and biosynthesis saturation level (*γ*_*B*_(*ϕ*_*B*_)). As the biosynthesis fraction increases, there are fewer provisioning enzymes to supply each biosynthetic enzyme, and the biosynthesis saturation level decreases. Figure 3a shows a graphical representation of this trade-off. To quantify the trade-off and find the optimal allocation, we choose the following function that approximates biosynthesis saturation level (*γ*_*B*_(*ϕ*_*B*_)) using the ratio of substrate supply (*M*_*Total*_(*ϕ*_*P*_ *ϵ*_*P*_ − *S*_*M*_ + *c* −*D*)) and maximum rate of macromolecule production (*M*_*Total*_(*ϕ*_*B*_ *ϵ*_*B*_ − *D*))(notice that *M*_*Total*_ cancels out):

**Figure 3.**
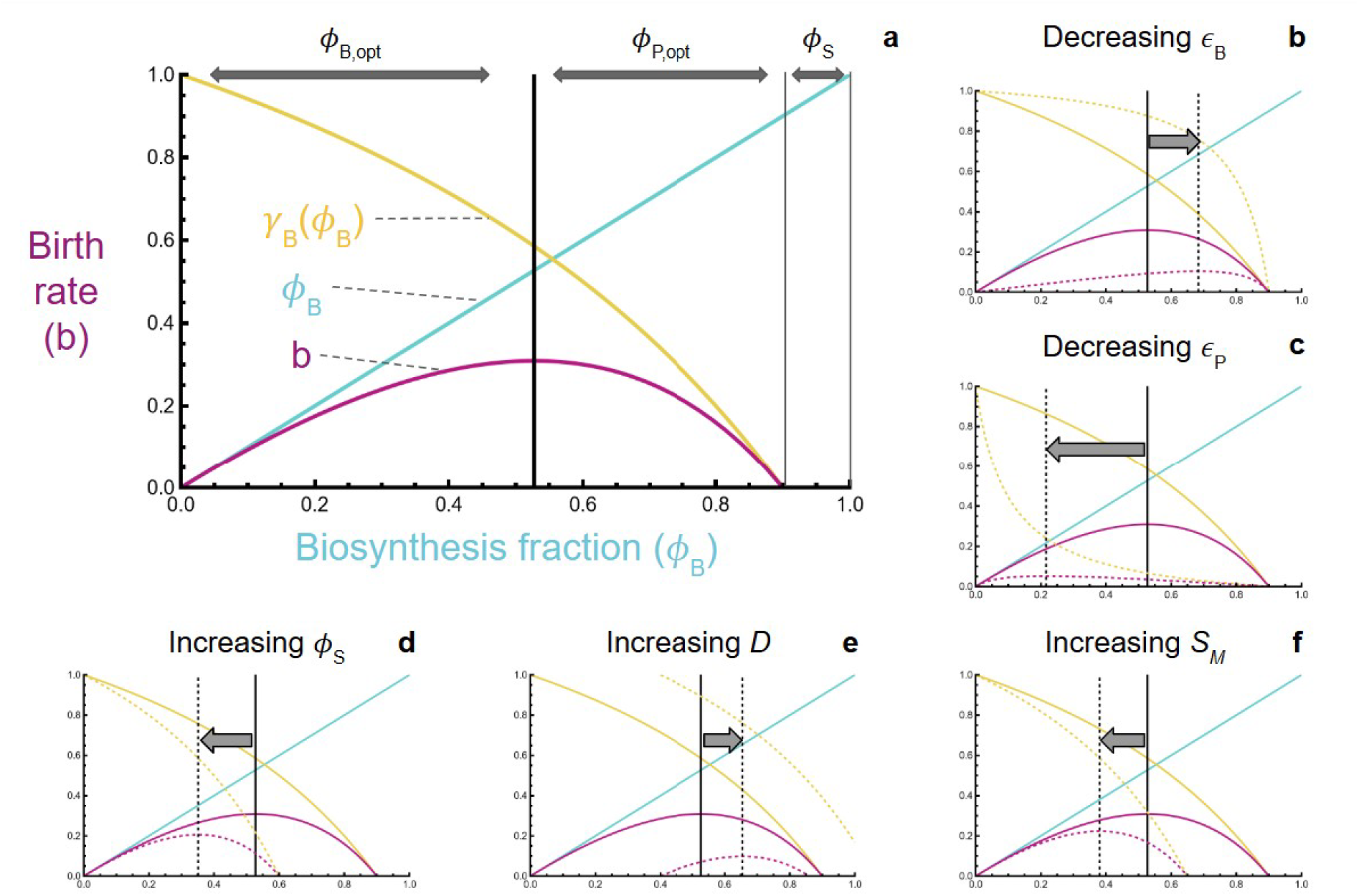
a) As modeled in equation (1), the birth rate (*b*, purple humped curve) is the product of the biosynthesis fraction (*ϕ*_*B*_ along the x-axis, blue line) and the speed of biosynthetic machinery (*ϵ*_*B*_ *γ*_*B*_(*ϕ*_*B*_), yellow declining curve) minus the degradation rate. When the provisioning fraction is maximal (biosynthesis fraction near zero), the speed of biosynthetic machinery is maximal (*ϵ*_*B*_), but decreases as macromolecules are shifted from provisioning (*ϕ*_*P*_) to biosynthesis (*ϕ*_*B*_). This shift leads to the birth rate rising, however, until an optimal allocation is reached (vertical black bars). In panels b-f, dashed lines show the trade-off after varying one of the parameters, while solid lines represent the baseline (shown in panel a). Decreasing the biosynthesis efficiency (*ϵ*_*B*_ in panel b) increases the optimal biosynthesis fraction, however, decreasing the provisioning efficiency (*ϵ*_*P*_ in panel c) has the opposite effect. Increasing the structural fraction (*ϕ*_*S*_ in panel d) decreases the provisioning and biosynthesis fractions. Increasing the degradation rate (*D* in panel e) increases the optimal biosynthesis fraction (given that recycling is efficient), however, increasing maintenance demands (*S*_*M*_ in panel f) decreases the optimal biosynthesis fraction. The following parameter values were used to obtain the solid lines: *ϵ*_*B*_ = 1, *ϵ*_*P*_ = 2, *ϕ*_*S*_ = 0.1, *D* = 0, *S*_*M*_ = 0, and *c* = 0.8. Dashed lines in panels b-f differ in the following parameter values: panel b: *ϵ*_*B*_ = 0.2; panel c: *ϵ*_*P*_ = 0.1; panel d: *ϕ*_*S*_ = 0.4; panel e: *D* = 0.4; panel f: *S*_*M*_ = 0.5.

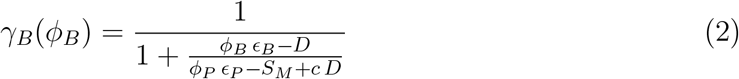

When biosynthesis is the limiting process and the substrates are supplied into the substrate pool at a rate much faster than the maximum rate of macromolecule production (*ϕ*_*P*_ *ϵ*_*P*_ − *S*_*M*_ + *c D >> ϕ*_*B*_ *ϵ*_*B*_ − *D*), the ratio of substrates to macromolecules in the cell is high, and the biosynthetic enzymes become saturated (*γ*_*B*_(*ϕ*_*B*_) = 1). On the other hand, when provisioning is the limiting process and the supply of substrates (including energy) is significantly smaller than the maximum rate of macromolecule production (*ϕ*_*P*_ *ϵ*_*P*_ −*S*_*M*_ + *c D << ϕ*_*B*_ *ϵ*_*B*_ −*D*), the ratio of substrates to macromolecules in the cell is low and the saturation level of biosynthetic enzymes approaches zero. Equation (2) is one of the main differences between our model and Scott et al. (2010), which assumes the saturation level of ribosomes is constant. In S4 Text, we explicitly model substrate dynamics and discuss the accuracy of equation (2).

## Results

### Optimal allocation

Shifting the allocation of macromolecules between provisioning (*ϕ*_*P*_) and biosynthesis (*ϕ*_*B*_), while holding the structural fraction constant (*ϕ*_*P*_ + *ϕ*_*B*_ = 1 − *ϕ*_*S*_), cell birth rate is maximized at a single optimal allocation (Figure 3a), with selection favoring genetic or plastic changes that bring the cell closer to the optimum. To determine the optimal allocation, we find the biosynthesis fraction (*ϕ*_*B*_) where the derivative of the birth rate equation (1) with respect to *ϕ*_*B*_ equals 0. The optimal biosynthesis fraction thus equals:

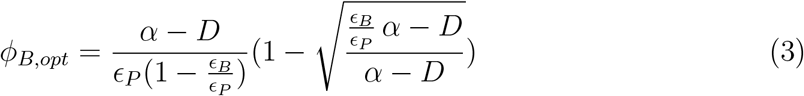

where *α* = *ϵ*_*P*_ (1 −*ϕ*_*S*_) −*S*_*M*_ + *c D* is the maximum substrate supply rate (when the provisioning fraction is maximal), which must be positive. To better understand this result, let us first focus on when degradation and maintenance demands are absent (*D* = *S*_*M*_ = 0). Under this condition, the optimal biosynthesis fraction simplifies to 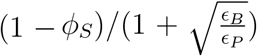, and the optimal provisioning fraction (which equals 1 − *ϕ*_*S*_ − *ϕ*_*B,opt*_) simplifies to 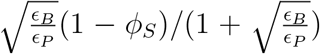. The ratio of the provisioning fraction over the biosynthesis fraction is thus inversely related to the ratio of provisioning over biosynthesis efficiency and equals:

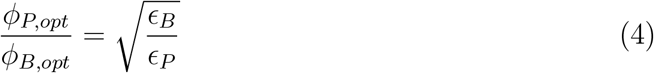

This shows that if, for example, the biosynthesis fraction is two times more efficient than the provisioning fraction 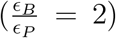, the provisioning fraction will be larger than the biosynthesis fraction but not large enough to completely compensate for the lower efficiency 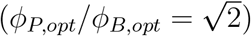, therefore the less efficient pathway remains more limiting. This highlights that optimal allocation is determined by a balance between the production of more efficient macromolecules and reducing the degree of limitation imposed by the less efficient pathway on macromolecule production.

When the degradation rate is low, the optimal allocation can be approximated by equations (5) and (6) to leading order in *D* (using first-order Taylor series expansion), which we use to guide our understanding of the impacts of maintenance demands (*S*_*M*_), degradation (*D*), and recycling efficiency (*c*) (see Figure 3e and 3f).

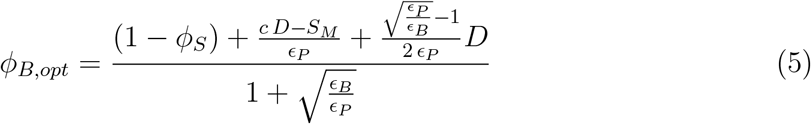

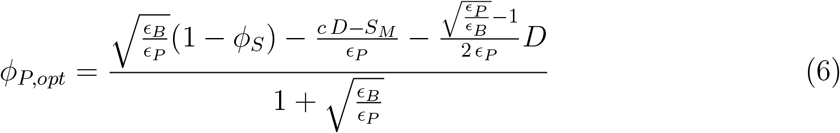

The optimal provisioning fraction increases with increasing maintenance demands (*S*_*M*_) or decreasing recycling (*c D*) to compensate for the lower substrate supply rate (Figures 3e and 3f). Notice that the larger the provisioning efficiency, the smaller the effect of recycling and maintenance demands on the optimal provisioning and biosynthesis fractions (see 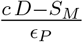 in the numerator of equations (5) and (6)). This is because when provisioning machinery is fast, small changes in the provisioning fraction can compensate for the effect of recycling and maintenance demands on substrate supply.

Degradation affects the optimal allocation even when there is no recycling (see 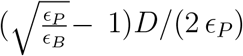 in the numerator of equations (5) and (6)). This is because increasing the degradation rate reduces the birth rate of cells, thereby reducing substrate supply to new cells at a rate equal to 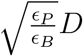, which is proportional to steady-state substrate concentration when degradation rate is low (equation S5). Degradation also acts as a sink for substrates at a rate equal to *D*. Therefore, increasing the degradation rate increases the optimal biosynthesis fraction when biosynthesis is more limiting 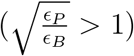 and has the opposite effect when provisioning is more limiting 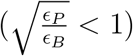.

The saturation level of biosynthesis under optimal allocation can be calculated by plugging in the optimal biosynthesis fraction (equation (3)) into equation (2):

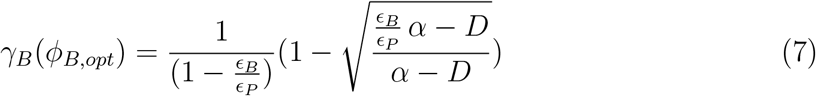

Under optimal allocation and in the absence of degradation (*D* = 0), biosynthesis saturation level simplifies to 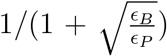, which is maximal (*γ*_*B*_(*ϕ*_*B,opt*_) = 1) when biosynthesis is limiting (*ϵ*_*P*_ *>> ϵ*_*B*_), and approaches zero when provisioning is limiting (*ϵ*_*B*_ *>> ϵ*_*P*_).

To determine the effect of degradation on the biosynthesis saturation level, we investigate its derivative with respect to *D* which equals 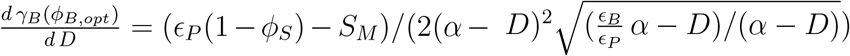. The denominator of this fraction must be positive, and the numerator must also be positive if the birth rate is positive (otherwise substrate loss is larger than the maximum substrate provisioning); therefore, degradation always increases biosynthesis saturation level. This is because the degradation rate reduces the reproduction rate and, consequently, the substrate supply to new cells, which in turn increases the concentration of substrates in the cytoplasm and the biosynthesis saturation level. The effect of maintenance demands on biosynthesis saturation level depends on the degradation rate 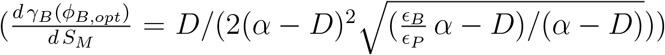. Interestingly, in the absence of degradation, maintenance demands do not affect the biosynthesis saturation level. This is because the effect of maintenance demands on the biosynthesis and provisioning fractions compensates for its impact on substrate supply.

To calculate the birth rate under optimal allocation (*b*_*opt*_), we plug in *ϕ*_*B,opt*_ (equation (3)) and *γ*_*B*_(*ϕ*_*B,opt*_) (equation (7)) into equation (1).To facilitate interpretation, the resulting expression can be approximated using a first-order Taylor series expansion with respect to the degradation rate, assuming the degradation rate is low:

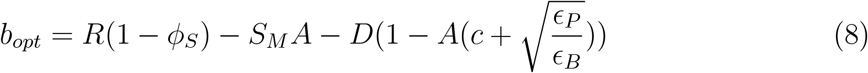

where *R* (the return on investment into growth) equals 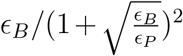, and *A* (measuring the extent to which provisioning is limiting) equals 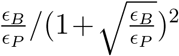. Equation (8) indicates that the birth rate equals the fraction of macromolecules allocated to provisioning and biosynthesis (1 − *ϕ*_*S*_ = *ϕ*_*B*_ + *ϕ*_*P*_) times the return on investment (*R*) minus the loss due to maintenance demands (*S*_*M*_ *A*) and degradation 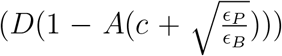. The return on investment (*R*) is related to the harmonic mean of the provisioning and biosynthesis efficiencies 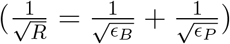 and is closer to the efficiency of the less efficient process.

The effect of degradation and maintenance demands on the birth rate depends on which pathway is limiting. When biosynthesis is limiting (*ϵ*_*P*_ *>> ϵ*_*B*_, *A* ≈ 0), the reduction in the birth rate due to degradation is maximal and maintenance demands have almost no effect on the birth rate. On the other hand, when provisioning is limiting growth (*ϵ*_*P*_ *<< ϵ*_*B*_, *A* ≈ 1), the effect of maintenance demands on the birth rate is maximal, while degradation’s effect depends on recycling efficiency. When the substrates are fully recycled after degradation (*c* ≈ 1), the cost of degradation is minimal, however, when the substrates are not recycled (*c* ≈ 0), the cost of degradation is high (Figure S3).

### Cost of traits

For a trait conferring a given benefit (e.g., a given reduction in death rate), we provide a framework to investigate the cost of the trait. We also explore how this cost will be affected by provisioning and biosynthesis efficiency, which are determined by environmental quality and genetic background (Figure 4).

**Figure 4.**
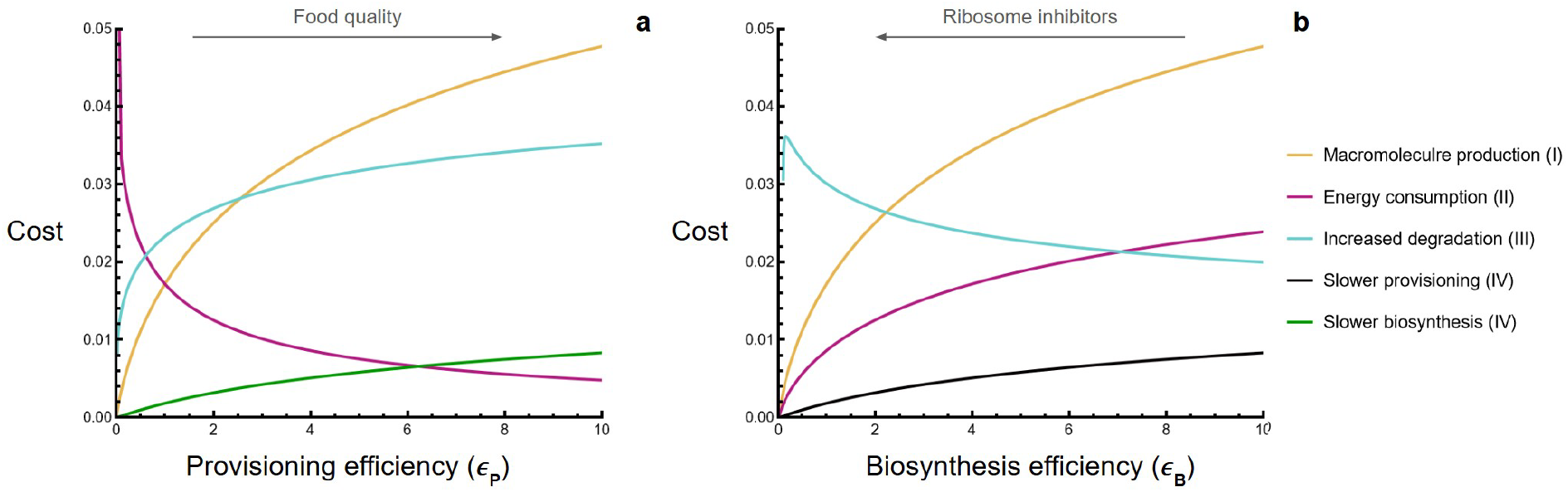
The effect of altering provisioning efficiency (e.g., by food quality, panel a) and biosynthesis efficiency (e.g., by antibiotics that inhibit ribosomes, panel b) on the cost of different categories of traits (points I-IV). The cost is calculated as the absolute difference between the birth rate of a reference cell and a cell possessing the trait. The possession of the trait alters one of the parameter values: macromolecule production increases *ϕ*_*S*_, energy consumption increases *S*_*M*_, increased degradation increases *D*, slower provisioning decreases *ϵ*_*P*_, and slower biosynthesis decreases *ϵ*_*B*_, all by 0.05. The parameter values for the reference cell are *ϕ*_*S*_ = 0.5, *c* = 0.8, *D* = 0, *S*_*M*_ = 0, *ϵ*_*B*_ = 2 (panel a), and *ϵ*_*P*_ = 2 (panel b). The exact equations (3) and (7) were used to calculate the birth rate (not the Taylor series approximation).

Using equation (8), we investigate the cost of four different examples of traits conferring stress-tolerance: I. producing a macromolecule that mitigates stress (e.g., an enzyme degrading a toxin or a chelator neutralizing it). II. increasing energy consumption for maintenance (e.g., energy allocation for higher activity of preexisting pumps). III. increasing macromolecule degradation (e.g., to reduce the accumulation of damage in the cell). IV. using a slower metabolic pathway that reduces the cell’s vulnerability to stress.

1. *Structural macromolecule allocation*-Allocating macromolecules for stress tolerance increases the structural fraction, therefore the change in birth rate equals −Δ*ϕ*_*S*_ *R*, where delta (Δ) symbolizes the change in a parameter. The lower the return on investment, the less costly the trait. In other words, the cost of allocating macromolecules to a trait depends on the growth that could have been achieved if those macromolecules were allocated to provisioning and biosynthesis. When the return on investment is higher, there is more to lose.
2. *Higher energy allocation to maintenance*-Increasing the activity of preexisting pumps requires energy allocation and increases the maintenance demands, thus the change in birth rate equals −Δ*S*_*M*_ *A*. The increase in maintenance demand is more costly when provisioning is limiting (*A* ≈ 1). Any cost involving the allocation of energy or substrates (e.g., activating efflux pumps, feeding symbionts, and secreting metabolites) follows the same pattern.
3. *Increased degradation rate*-The change in the birth rate due to an increase in the degradation rate depends on the recycling efficiency and the extent to which provisioning is limiting and equals 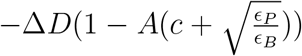. If recycling is efficient (*c* ≈ 1), the cost of degradation is maximal when biosynthesis is limiting (*A* ≈ 0). However, if recycling is inefficient (*c* ≈ 0), the cost of degradation is overall high, and is relatively independent of provisioning and biosynthesis efficiencies (Figure S3).
4. *Switching to slower pathways*-Switching to a slower metabolic pathway reduces the provisioning, biosynthesis efficiency, or both, thereby lowering the return on investment (*R*) and altering the extent to which provisioning is limiting (*A*). The change in birth rate will thus be 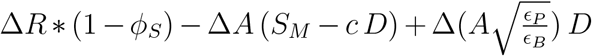. The more limiting a fraction is, the more detrimental reducing its efficiency would be on *R*. Therefore, as biosynthesis efficiency increases, the cost of slowing provisioning increases, and vice versa (Figure 4). Increasing the structural fraction reduces the provisioning and biosynthesis fractions, thus decreasing the cost of slowing down provisioning or biosynthesis. The cost of slowing down provisioning (Δ*A >* 0) increases with maintenance demands and decreases with recycling. The opposite is true for the cost of slowing down biosynthesis (Δ*A <* 0). Degradation has an additional effect on the cost of slowing down provisioning or biosynthesis, which depends on which pathway is more limiting 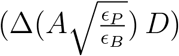.

This framework thus helps predict the relative costs of different types of traits, in addition to how these costs depend on environmental conditions and genetic background, a point to which we return in the Discussion.

### Bacterial growth laws

We next examine the correspondence between this model and the bacterial growth laws (Figure 1a). We assume cells have reached optimal allocation and focus on the relationship between birth rate (*b*_*opt*_), biosynthesis fraction (*ϕ*_*B,opt*_), and biosynthesis saturation (*γ*_*B*_(*ϕ*_*B,opt*_)) (Figure 5). As the death rate is small during exponential growth (0.001 −0.05 *h*^− 1^ depending on the age of the cells[25]), the birth rate can be a proxy for the growth rate under the laboratory conditions used to study growth laws. Moreover, biosynthesis fraction and biosynthesis saturation level are correlated with the ribosome fraction in the proteome and the average speed of ribosomes (which equals elongation rate times the fraction of active ribosomes), respectively, which we use as proxies. We consider manipulating provisioning efficiency here (e.g., via foods with different quality) and consider manipulating biosynthesis efficiency in S3 Text (e.g., via ribosome inhibitors).

**Figure 5.**
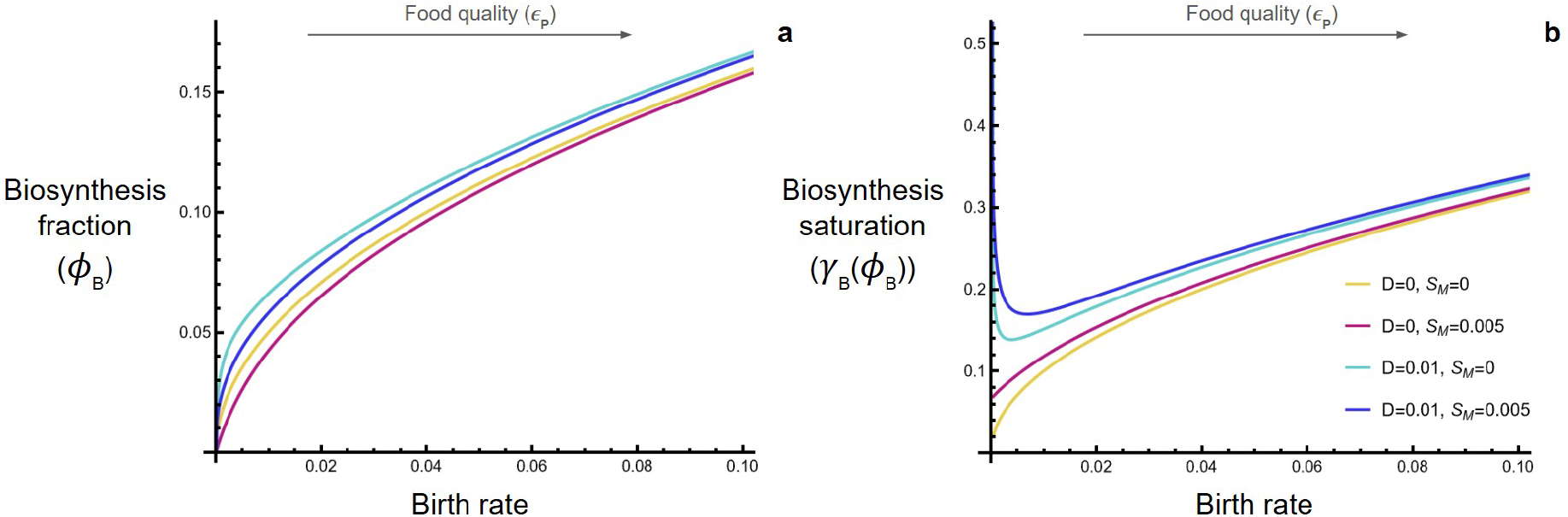
Degradation, recycling, and maintenance demands affect the shape of bacterial growth laws, particularly during slow growth. To compare the model predictions with the growth laws, the biosynthesis fraction (panel a) is used as a proxy for the ribosome fraction (Figure 1b) and the biosynthesis saturation level (panel b) as a proxy for the average speed of ribosomes (Figure 1c). The exact equations (3) and (7) were used to calculate the birth rate, optimal biosynthesis fraction, and biosynthesis saturation (not the Taylor series approximation). Notice that during fast growth, the slope of all lines in panel a is similar to the slope of the red line and approximately equals 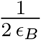. Other parameter values: *ϕ*_*S*_ = 0.5, *ϵ*_*B*_ = 2, and *c* = 0.8.

In the absence of degradation and maintenance demands (*D* = *S*_*M*_ = 0), as provisioning efficiency is increased, the birth rate, biosynthesis fraction, and the biosynthesis saturation level all increase (Figures 5a and 5b, yellow lines). Because the biosynthesis saturation level is variable, the relationship between birth rate and biosynthesis fraction is nonlinear, particularly during slow growth (Figure 5a, yellow line, whose slope is given by 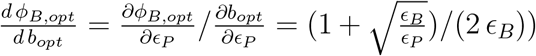. During fast growth, the biosynthesis fraction is saturated (because *ϵ*_*P*_ *>> ϵ*_*B*_), therefore the slope approaches a constant value of 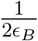. During slow growth, however, the biosynthesis fraction and saturation level approach zero, contradicting empirical observations (Figure 1a).

Allowing for degradation and maintenance demands leads to a broad range of predictions as growth slows. With only maintenance demands (*S*_*M*_ *>* 0, *D* = 0), as the birth rate approaches zero, the biosynthesis saturation level approaches a non-zero value 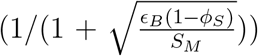, in keeping with observed patterns (Figure 1c); the biosynthesis fraction continues to approach zero, however (Figures 5a and 5b, red lines).

In the presence of degradation (*D >* 0, regardless of *S*_*M*_), the biosynthesis fraction and the biosynthesis saturation level approach non-zero values as growth rate slows to zero (with intercepts of 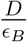 and 1, respectively). The shape of these curves at low birth rates does not, however, match observations (compare Figures 5a and 5b, dark and light blue curves, to Figures 1a and 1c, respectively), with the predicted biosynthesis fraction being concave-down at low birth rates, while the predicted biosynthesis saturation level is concave-up. More empirical measurements at low growth rates would help determine whether these discrepancies are real, given that previous studies have not investigated very low growth rates (e.g., the smallest growth rate investigated by Dai et al. (2016) is 0.03 *h*^− 1^). We return to the behavior of the growth laws at low growth in the Discussion. In the supplementary material, we motivate our choice of parameters (S2 Text), explore the parameter space more thoroughly (Figures S5 and S6), and investigate an alternative model that explicitly tracks substrate concentration and yields qualitatively similar results (S4 Text).

## Discussion

The trade-off between allocation to biosynthesis and provisioning is a universal optimization problem all organisms face. By modeling this trade-off generally, the model we explore transitions smoothly between different factors limiting growth. This model can predict the fitness surface, i.e., the birth rate under different allocations (Figure 3), and the growth of an organism not at the optimum, as well as predicting optimal allocation patterns (Figures 4 and 5). Using this model, we predict the effect of environmental change and genetic background on the cost of various traits (Figure 4). Notably, we predict that the cost of traits that involve the production of a new macromolecule depends on the overall speed of provisioning and biosynthesis (point I). On the other hand, the cost of traits that require energy allocation (point II) or increased degradation (point III) depends on how limiting provisioning and biosynthesis are. Furthermore, we showed that the energetic demands of maintenance, degradation, and recycling affect the shape of bacterial growth laws, particularly during slow growth (Figure 5). Overall, we argue that combining provisioning and biosynthetic limitations provides a broader framework to think about the costs of traits than just focusing on energy or ribosome allocation.

We focus on the allocation of all macromolecules instead of just proteins and suggest that the relationship between ribosome fraction and growth rate is a specific case of the more general relationship between the biosynthesis fraction and growth rate. This relationship can be explained without assuming that protein production or any other specific process is the limiting step in growth. This contrasts with previous studies that focused on ribosome allocation and argued that protein production is the main limiting step in growth due to its high energetic cost, the large fraction of proteins in the microbial dry mass, and the self-replicating nature of ribosomes. [23, 15, 6].

While our model provides a unified framework to understand the costs of different traits and patterns observed (the bacterial growth laws) when growing rapidly, our model does not reproduce the shape of growth laws during slow growth (Figure 5). This discrepancy suggests that additional processes are playing an important role. For example, it has been suggested that the degradation rate increases during slow growth, which could explain the concave-up shape of the growth laws during slow growth[5](but a more recent study disagrees [10]). Alternatively, cells may regulate ribosome fractions in a manner that is non-optimal[26]. It has also been suggested that cells have evolved to grow in a fluctuating environment and maintain more ribosomes than needed during starvation, which is beneficial upon increased food availability[7, 18]. Our model supports the last explanation by showing that when food quality is very low, having a higher than optimal biosynthesis fraction results in a relatively small reduction in birth rate (i.e., the fitness surface is asymmetric; Figure 3c). Moreover, the regulation of gene expression is noisy in bacteria, and there is considerable heterogeneity in the allocation of individual cells [1]. Therefore, selection acting on an asymmetric fitness surface (Figure 3c, provisioning is limiting) may have resulted in a higher than optimal average biosynthesis fraction in the population. Nevertheless, a recent study has argued that previous empirical studies might have overestimated the ribosome fraction during slow growth by not giving cells enough time to reach steady state before measurement[6], casting doubt on whether growth laws are concave-up during slow growth.

Our model can predict the effect of environmental quality on the type of traits more likely to evolve (Points I-IV, Figure 4). All else being equal, a trait is more likely to evolve under conditions where it is less costly. Our model predicts that the cost of structural traits that require macromolecule allocation is the lowest in low-quality environments, where growth is already slow due to low food quality or slow biosynthesis (point I). On the other hand, the cost of energy allocation (which increases maintenance demands) is lowest when biosynthesis is limiting (due to high food quality or slow biosynthesis, point II). Alternatively, the cost of degradation is minimal when recycling is efficient and provisioning is limiting (due to low food quality or fast biosynthesis, point III). Lastly, the cost of slowing down provisioning (or biosynthesis) is the least when it is not limiting and the structural fraction is large (point IV). Moreover, our model can make qualitative predictions about the likelihood of stress tolerance evolution in a specific environment based on the types of possible mutations. For example, if becoming resistant to a stressor is only possible by traits that require energy allocation, we predict resistance to evolve more readily in environments where food quality is high or biosynthesis is inhibited. Notice that we quantify cost as an absolute reduction in the birth rate of cells caused by the trait, which is appropriate for continuously growing cells.

Our work highlights that to understand the effect of environmental quality on the cost of traits, multiple dimensions of environmental quality should be considered simultaneously. For example, although decreasing food quality (i.e., provisioning efficiency) increases the cost of energy-consuming traits, inhibiting biosynthesis (e.g., by antibiotics) has the opposite effect (point II). The same applies to the effect of genetic background on the cost of traits. A recent study did not find any meaningful relationship between the cost of traits and environmental quality, which may be because the environments differed in many factors with confounding effects on provisioning and biosynthesis efficiency[11]. We recommend that future studies determine to what degree provisioning and biosynthesis are limiting growth in different environments (using RNA-to-protein ratio as a potential proxy), and compare the cost of traits in similar environments that only differ in provisioning or biosynthesis efficiency. While these efficiencies are abstract quantities that cannot be directly measured, they can be inferred from the slope of the relationship between the biosynthesis fraction and the birth rate during fast growth (e.g., a slope of 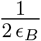 in Figure 5a).

Biomass production is a fundamental process required for the growth and reproduction of all living organisms. Our model may thus shed light on the trade-off between allocation to provisioning, biosynthesis, and structure at other levels of organization. For example, the prevalence of structural adaptations in desert-living plants (e.g., thorns) might be partially explained by the lower cost of structural traits in low-quality environments (or life-histories) for growth (Point I). Wherever biomass production is maximized, insights from this model might be applicable.

## Supporting information

S1 Text. **Assumptions and limitations**.

S2 Text. **Estimating parameter values**.

S3 Text. **Growth laws when partially inhibiting biosynthesis**.

S4 Text. **Explicitly modelling substrate dynamics**

Mathematica notebook. **Derivations and code for reproducing graphs**.

## Acknowledgments

We would like to thank David Anderson, Dan Kehila, Mary O’Connor, Steven Frank, Penelope Kahn, Anna Bazzicalupo, Eully Ao, Alireza Tafreshi, Caleigh Charlabois, Gabriel Greenberg-Pines, William Ou, Jawad Sakarchi, Austin Burt, Matthew Scott, Christian Landry, Laura Parfrey, Nobuhiko Tokuriki, Katie Marshall, Christopher Klausmeier, Elena Litchman, Maria Rebolleda-Gomez, Erol Akçay, Sara Mitri, Charles Mullon, Prompt Suathim, Amy Forsythe, Asad Hassan, Bryn Whiley, Nicola Love, Margaret Slein, Nicole Knight, Keila Stark, Kaleigh Davis, Fruin Pow, and many members of the Biodiversity Research Center for discussions and feedback. This research was supported by the Natural Sciences and Engineering Research Council of Canada Discovery Grant to S.P.O. (RGPIN-2022-03726).

## Notes

### Competing Interest Statement

The authors have declared no competing interest.

