## Supplementary information for "Bridging energy and ribosomal allocation models to predict the cost of traits in different environments"

#### Contents

1 Assumptions and limitations 2

2 Estimating parameter values 2

3 Growth laws when partially inhibiting biosynthesis 3

4 Explicitly modeling substrate dynamics 4

### 1 Assumptions and limitations

We assumed cells are under steady state, and biomass is continuously growing. We did not explicitly consider cell division events and assumed their energetic cost, i.e., the energy required for the reorganization of macromolecules and regulatory processes that accompany cell division, is part of maintenance demands. We assumed the maintenance demands are independent of the birth rate, however, because the energetic cost of cell division is considered part of maintenance demands, maintenance demands should increase with the birth rate. The same argument holds for the degradation rate.

We assumed that $\epsilon_P$, $\epsilon_B$, $S_M$, $D$, and $c$ are independent of the allocation. This is not entirely accurate because the size of fractions affects cell stoichiometric compositions[5], thereby potentially altering all the parameters. Accounting for this can explain some observations. For example, a study observed that cells under phosphate limitation reach a similar growth rate to cells under carbon and nitrogen limitations but with fewer ribosomes [6]. The lower allocation to the biosynthesis fraction in cells under phosphate limitations is consistent with optimal allocation if we consider that increasing the biosynthesis fraction increases the cell's phosphate content (because the biosynthesis fraction is rich in RNA), which decreases the provisioning efficiency when phosphate is scarce. Furthermore, a constant provisioning efficiency means that food uptake rises linearly with the cell's investment in the provisioning fraction, and there are no diminishing returns. This assumption is more accurate in well-mixed and nutrient-rich environments where diffusion of food into the cell's membrane is not limiting.

We also assumed provisioning efficiency is independent of maintenance demands. As maintenance mainly consumes energy, increasing maintenance demands increases energy production compared to other substrates. This increases the subfraction of energy production machinery in the provisioning fraction, which can alter provisioning efficiency. Similarly, because different macromolecules have different degradation rates, increasing the degradation rate can alter the ratio of biosynthetic machinery allocated to the production of each category of macromolecules, which can affect biosynthesis efficiency. We also assumed both degradation and maintenance demands are independent of the macromolecular composition of the cell and scale linearly with cell mass, which is not completely accurate. Moreover, we assumed that all the substrates used by the biosynthesis machinery are converted to macromolecules and there is no mass loss.

### 2 Estimating parameter values

As more than half of E.coli biomass is made of proteins[11], we estimate most of the parameters by just focusing on proteins:

**Degradation rate ($D$)**: According to Pine (1973), the average degradation rate of proteins in E.coli is about $0.01\,hour^{-1}$ and increases to about 0.04 as the growth rate slows[9]. However, Gupta et al. (2024) have found that protein degradation rates are generally independent of growth rate. This study found that the degradation rates are higher in the N-limitation regime (around $0.02 - 0.04\,hour^{-1}$) compared to the C-limitation and the P-limitation (around $0.01\,hour^{-1}$)[4].

**Maintenance demands ($S_M$)**: According to Lynch and Marinov (2015), the energetic cost of cell maintenance ($C_M$, unit: $(10^9 ATP)$/(cell*hour)) scales with cell volume ($V$, unit: $\mu m^3$) as follows:$C_M = 0.39V^{0.88}$[7]. *E. Coli*'s volume is roughly$1\,\mu m^3$ and its mass is roughly $1\,pg$ . Therefore $C_M = 0.39 * 1 * (10^9 ATP)/(cell * hour)$. To convert

$C_M$ to $S_M$, we assume $3 * 10^1$ ATP molecules are produced per Glucose molecule during aerobic respiration. $S_M$ roughly equals the mass of glucose used for maintenance per unit mass of macromolecules (which is roughly equal to cell mass), per unit time (molar mass of Glucose $=1.8 * 10^2\, daltons$):

$$S_M = 0.39 * 10^9 * \frac{1}{30} * \frac{180}{(6.022 * 10^23)} * \frac{1}{10^{-12}} = 4 * 10^{-3}\, hour^{-1}$$

This probably underestimates the true value of $S_M$, because $S_M$ also incorporates the metabolites that are secreted by the cells.

**Biosynthesis efficiency ($\epsilon_B$)**: Scott et al (2010). has estimated translational capacity ($\kappa_n$, which is similar to $\epsilon_B$ in our model) for a few strains of *E. coli* in the range 1.4 to 4.0. According to Metzl-Raz (2017), the slope of the relationship between ribosome fraction and growth rate in *S. cerevisiae* is $\frac{0.35}{ln(2)} = 0.5$[8]. Our model predicts that the slope should be about $\frac{1}{2\,\epsilon_B}$. Therefore $\epsilon_B = 1\, hour^{-1}$.

**Structural fraction ($\phi_S$)**: Scott et al.(2010) has estimated the fixed fraction ($\phi_Q$) in the range 0.4-0.5 for a few strains of *E. coli* [10]. As the biosynthesis fraction contains most of the RNAs (approximately 20% of cell mass[11]) and the structural fraction also contains DNA, lipids, and carbohydrates (together around 10% of cell mass[11]), the structural fraction is roughly around 0.5.

**Recycling efficiency (c)**: The recycling efficiency must be less than 1 because the energy used in protein translation, folding, and localization is not recycled after degradation, and the degradation itself requires energy in Bacteria. Overall, most of the amino acids are recycled after degradation, however, we did not find any quantitative estimates. Therefore, we try to calculate a theoretical upper limit. Each amino acid requires one ATP for being loaded on a tRNA, in addition to two GTPs for being incorporated by a ribosome into a polypeptide. Therefore, roughly 1 glucose molecule (weight: $180 Daltons$) should be burned to provide energy ($3 * 10^1$ ATP) to incorporate the energy required for incorporating 10 amino acids (average weight: roughly$110 Daltons$). If all amino acids are recycled after degradation and the energetic cost of protein folding and localization is negligible, the recycling efficiency roughly equals 0.85, but it is likely smaller than that.

### 3 Growth laws when partially inhibiting biosynthesis

In the absence of degradation and maintenance demands, when biosynthesis efficiency is altered, there is a similar relationship between biosynthesis fraction ($\phi_{B,opt}$) and birth rate ($b_{opt}$) with a negative slope that equals $\frac{\frac{\partial \phi_{B,opt}}{\partial \epsilon_B}}{\frac{\partial b_{opt}}{\partial \epsilon_B}} = -\frac{1+\sqrt{\frac{\epsilon_P}{\epsilon_B}}}{2\,\epsilon_P}$ (Figure S5). During fast growth, where biosynthesis efficiency is larger than provisioning efficiency ($\epsilon_B >> \epsilon_P$), the slope approaches $\frac{-1}{2\epsilon_P}$. The presence of degradation and maintenance demands affects the relationship but does not change the overall pattern (Figure S1).

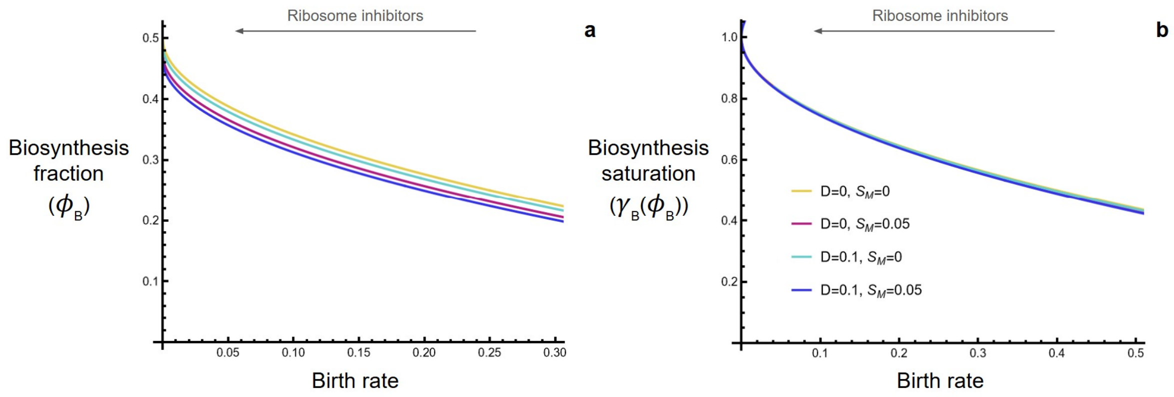


Figure S1: This figure shows the effect of degradation, recycling, and maintenance demands on the relationship between the birth rate, the biosynthesis fraction, and biosynthesis saturation level when altering biosynthesis efficiency (e.g., by partially inhibiting ribosomes). Under the optimal allocation, decreasing biosynthesis efficiency ($\epsilon_B$, from right to the left) decreases the birth rate ($b_{opt}$), while increasing the biosynthesis fraction ($\phi_{B,opt}$, panel a) and biosynthesis saturation level ($\gamma_B(\phi_{B,opt})$, panel b). Other parameter values: $\phi_S = 0.5$, $c = 0.8$, and $\epsilon_P = 2$

### 4 Explicitly modeling substrate dynamics

Here, we explicitly account for the mass concentration of substrates (S, hereafter concentration, dimension $= \frac{mass}{vollume}$) in the cell to investigate the accuracy of equation (2), which relates the saturation level of biosynthesis to biosynthesis fraction ($\gamma_B(\phi_B)$). The provisioning machinery, recycling, and maintenance demands supply substrates to the substrate pool at the rate $M_P\,\epsilon_P + M_{total}(c\,D - S_m)$ per cell. The substrate consumption rate by the biosynthesis fraction per cell equals $M_B\,\epsilon_B\,\gamma_B(S)$. We assume the saturation level of biosynthetic machinery is a hyperbolic function of substrate concentration, i.e., $\gamma_B(S) = \frac{S}{S+k_B}$, where $k_B$ is the half-saturation substrate concentration for the biosynthesis fraction (dimension: $\frac{mass}{volume}$). To calculate the effect of these fluxes on substrate concentration, we need to convert them from per cell to per unit volume by multiplying them by $\frac{\rho}{M_{Total}}$, where $M_{Total}$ is the mass of all cell macromolecules and $\rho$ is the density of macromolecules (which is slightly lower than biomass density because substrates are excluded, dimension: $\frac{mass}{volume}$). Finally, as the cell grows (at rate $b = \phi_B\,\epsilon_B \frac{S}{k_B+S} - D$, according to equation (1)), the substrates are diluted proportional to their concentration (S)[3]. Overall, we can write the change in concentration as follows:

$$\frac{dS}{dt} = (\underbrace{M_P\,\epsilon_P}_{\text{supply by }\phi_P} + \overbrace{M_{Total}\,c\,D}^{\text{recycling}} - \overbrace{M_{Total}\,S_M}^{\text{maintenance}} - \underbrace{M_B\,\epsilon_B \frac{S}{k_B+S}}_{\text{consumption by }\phi_B}) \overbrace{\frac{\rho}{M_{total}}}^{\text{conversion}} - \underbrace{(\phi_B\,\epsilon_B \frac{S}{k_B+S} - D)S}_{\text{dilution due to growth}} \tag{S1}$$

During balanced growth conditions, we can assume the substrate pool reaches a steady-state concentration ($\frac{dS}{dt} = 0$). This steady-state substrate concentration ($S_{ss}$)

equals:

$$S_{ss} = \rho \, \frac{\theta - \beta_{Max} + D \, \frac{k_B}{\rho} + \sqrt{(\theta - \beta_{Max} + D \, \frac{k_B}{\rho})^2 + 4 \, \theta \, \frac{k_B}{\rho} (\beta_{Max} - D)}}{2(\beta_{Max} - D)} \tag{S2}$$

where $\beta_{Max} = \phi_B \, \epsilon_B$ is the maximum substrate consumption rate by the biosynthesis fraction, and $\theta = \phi_P \, \epsilon_P - S_M + c \, D$ is the substrate supply rate. Therefore the saturation level of the biosynthesis fraction equals:

$$\gamma_B(S_{ss}) = \frac{1}{1 + \frac{k_B}{S_{ss}}} = \frac{1}{1 + \frac{2(\beta_{Max} - D) \frac{k_B}{\rho}}{\theta - \beta_{Max} + D \, \frac{k_B}{\rho} + \sqrt{(\theta - \beta_{Max} + D \, \frac{k_B}{\rho})^2 + 4 \, \theta \, \frac{k_B}{\rho} (\beta_{Max} - D)}}} \tag{S3}$$

As $k_B$ and $\rho$ always appear as $\frac{k_B}{\rho}$, only their ratio matters. When the density of macromolecules is higher, all reactions happen in a smaller volume, resulting in a higher concentration of substrates. Therefore $\frac{k_B}{\rho}$ is related to the ratio of the biosynthesis fraction substrate affinity to the substrate concentration and is negatively correlated with biosynthesis saturation level.

When $k_B$ is larger than $\rho$, equation (2) overestimates the speed of biosynthesis fraction and the birth rate, and when $\rho$ is larger than $k_B$, equation (2) underestimates the speed of biosynthesis fraction and birth rate (Figure S2). However, when $k_B = \rho$, equation (S3) simplifies to the approximation made by equation (2). Under this assumption, the steady state concentration of substrates simplifies to:

$$S_{ss} = \rho \, \frac{-D(1 + \sqrt{\frac{\epsilon_B}{\epsilon_P}})^2 + 2 \, \frac{\epsilon_B}{\epsilon_P} \, S_M - 2 \, \epsilon_B (1 - \phi_S)}{2 \, \frac{\epsilon_B}{\epsilon_P}^{\frac{3}{2}} (S_M - \epsilon_P (1 - \phi_S))} \tag{S4}$$

which can be approximated by a first-order Taylor series expansion with respect to degradation:

$$\rho \sqrt{\frac{\epsilon_P}{\epsilon_B}} + \frac{\rho \, (1 + \sqrt{\frac{\epsilon_B}{\epsilon_P}})^2}{2 \, (\frac{\epsilon_B}{\epsilon_P})^{\frac{3}{2}} (\epsilon_P (1 - \phi_S) - S_M)} \, D \tag{S5}$$

Because the biosynthesis fraction is an abstract entity, it is difficult to estimate $k_B$ accurately. Given that the concentration of substrates for most enzymes in *E. coli* is equal to or larger than the half-saturation concentration of their enzymes [1], and that the density of macromolecules in E. coli is roughly one order of magnitude larger than the mass concentration of metabolites [2, 1], we suspect that macromolecular density ($\rho$) should be at least one order of magnitude larger than $k_B$ in *E. coli* grown under normal conditions.

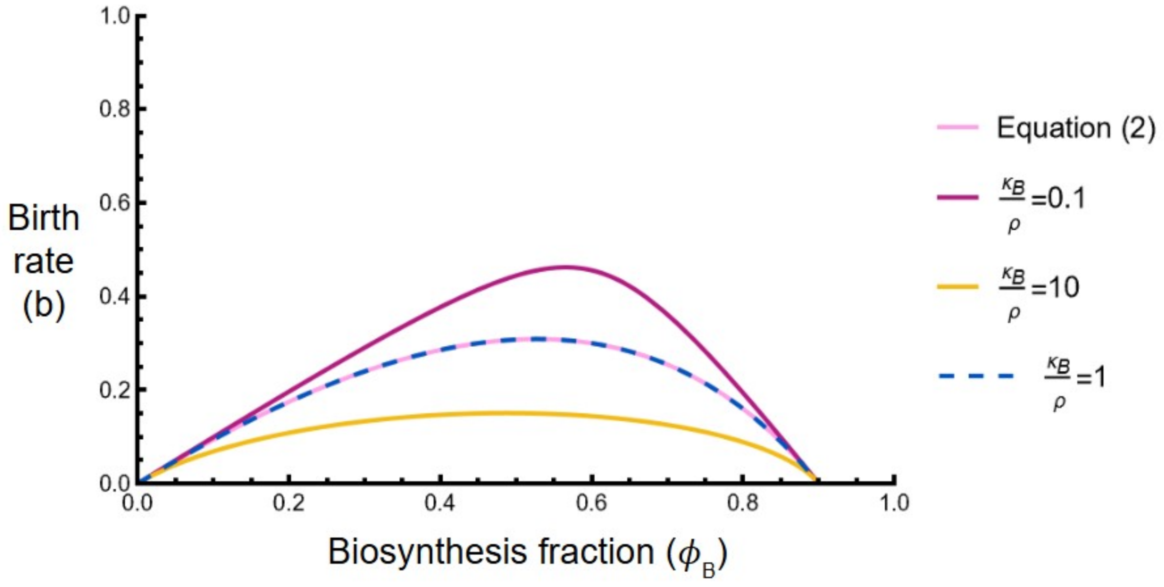


Figure S2: The accuracy of equation (2) (pink line) in approximating equation (S3) (obtained by explicitly modeling substrate dynamics) depends on the density of macromolecules ($\rho$) and half-saturation constant of biosynthesis fraction ($k_B$). When the density of macromolecules equals the half-saturation constant of biosynthesis fraction ($\rho = k_B$), the explicit solution for the speed of biosynthesis fraction equals the approximation made by equation (2) (dashed blue line). When $k_B$ is larger than $\rho$ equation (2) overestimates the speed of biosynthesis fraction and the birth rate (yellow line). When the $\rho$ is larger than $k_B$, equation (2) underestimates the speed of biosynthesis fraction and birth rate (purple line). Other parameter values: $\epsilon_B = 1$, $\epsilon_P = 2$, $\phi_S = 0.1$, $c = 0.8$, $D = 0$, and $S_M = 0$

Using equation (S3), we can find the optimal biosynthesis fraction (refer to the supplementary Mathematica file for analytical expressions) and then compare the results with the results obtained using equation (2). Figure S3 shows that the effect of the effect of the environment on the cost of traits is in qualitative agreement with predictions obtained using equation (2) (Figure 4). Figure S4 shows that the shape of bacterial growth laws is consistent when comparing the predictions obtained using equations (2) and (S3). Figures S5 and S6 show that the relationship between the biosynthesis fraction and birth rate (i.e., bacterial growth laws) obtained by equation S3 and equation (2), respectively, are always concave down during slow growth, however, Figure S5 is closer to being linear. Figure S7 shows the effect of changing the $\frac{k_B}{\rho}$ ratio on the shape of the bacterial growth laws.

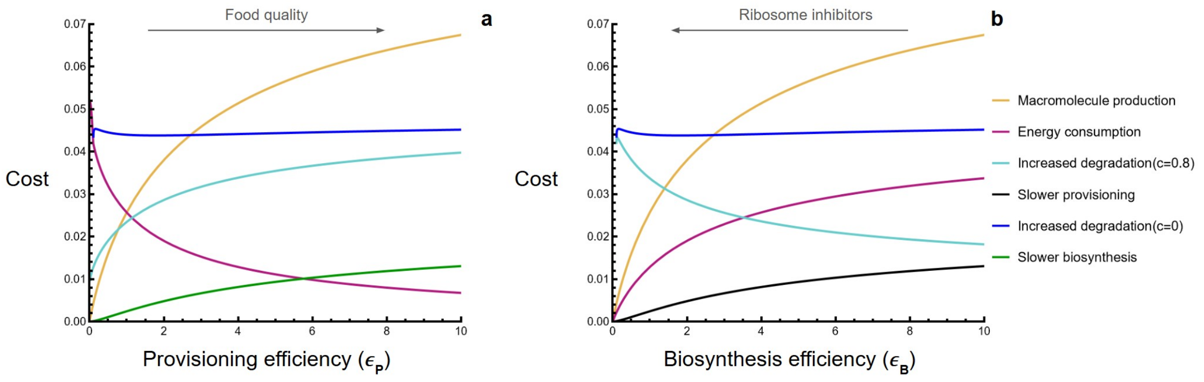


Figure S3: When using equation (S3), the effect of changing different parameters on the birth rate is in qualitative agreement with when equation (2) is used Figure 4). The panels show how the cost of traits is affected by changing provisioning efficiency ($\epsilon_P$ in panel a) and biosynthesis efficiency ($\epsilon_B$ in panel b). The cost is calculated as the absolute difference between the birth rate of a reference cell and a cell possessing the trait. The possession of the trait alters one of the parameter values: macromolecule production increases $\phi_S$, energy consumption increases $S_M$, increased degradation increases $D$, slower provisioning decreases $\epsilon_P$, and slower biosynthesis decreases $\epsilon_B$, all by 0.05. Parameter values for the reference cell: $\phi_S = 0.5$, $c = 0.8$, $D = 0$, $S_M = 0$, $k_B = 1$, $\rho = 10$, $\epsilon_B = 1$ (panel a), and $\epsilon_P = 2$ (panel b).

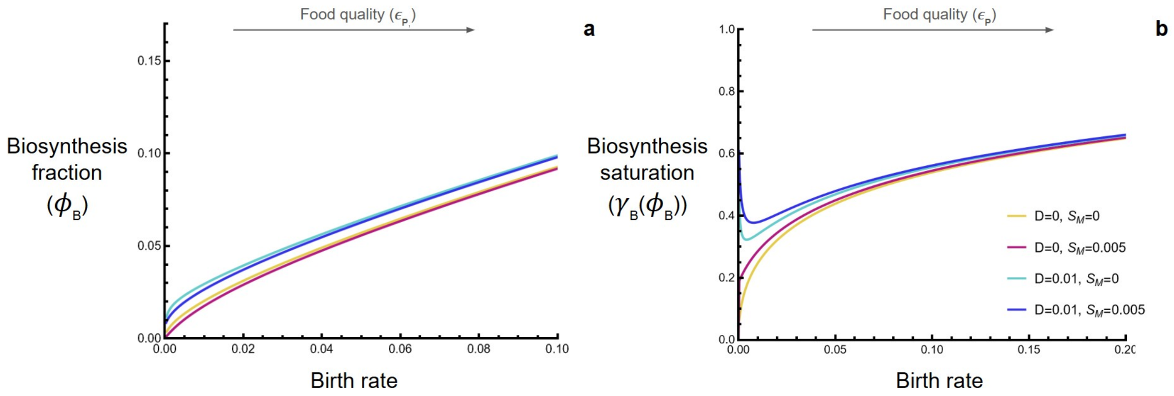


Figure S4: When using equation (S3), the predicted shape of bacterial growth laws, wen altering food quality ($\epsilon_P$), is in qualitative agreement with when equation (2) is used (Figure 5). Other parameter values: $\phi_S = 0.5$, $c = 0.8$, $\epsilon_B = 2$, $k_B = 1$, and $\rho = 10$

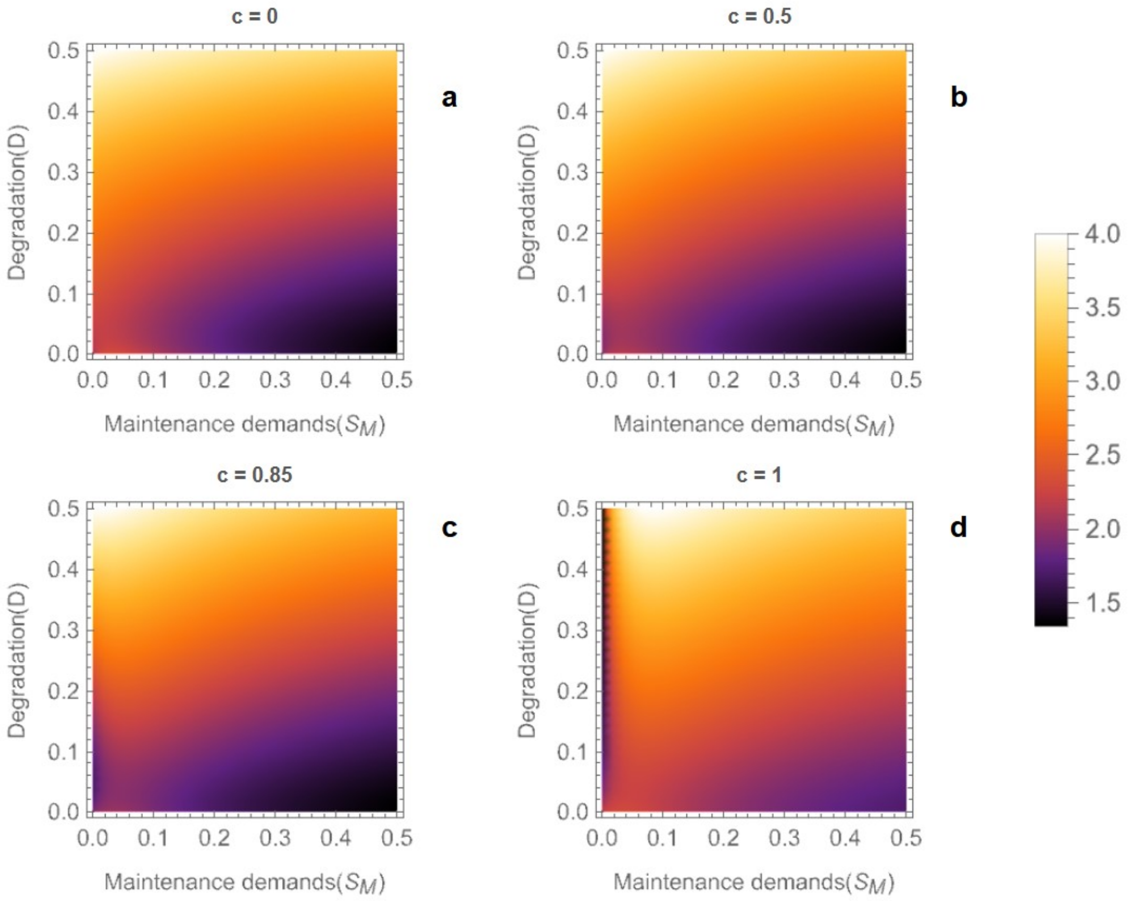


Figure S5: This figure explores the effect of degradation rate ($D$), recycling efficiency ($c$), and maintenance demands ($S_M$) on the direction of deviation of the growth laws from linearity during slow growth (when explicitly modeling substrate dynamics using equation (S3)). This deviation is shown in the heatmaps using the ratio of the slope of growth laws (Figure S4a) when the birth rate is low ($\epsilon_P$ is 0.1 higher than the value that results in a zero birth rate) over when the birth rate is high ($\epsilon_P = 20$). Different panels show different recycling efficiencies. If this ratio exceeds 1, the growth laws are concave down, and if the the ratio is between 0 and 1, the growth laws are concave up, similar to the observations. This figure shows that the ratio is always larger than one, meaning the growth laws are always concave down. Other parameter values: $\epsilon_B = 3$, $\phi_S = 0.5$, $\rho = 10$, and $k_B = 1$.

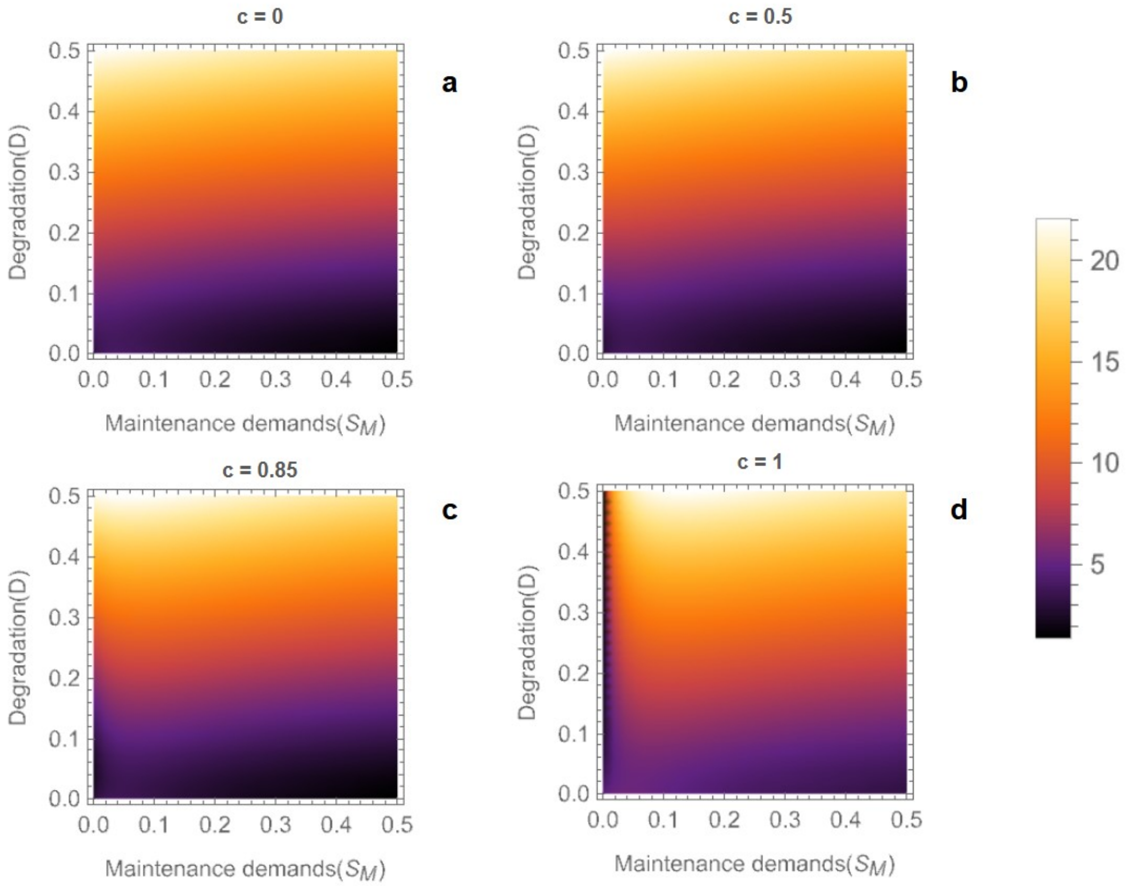


Figure S6: This figure explores the effect of degradation rate ($D$), recycling efficiency ($c$), and maintenance demands ($S_M$) on the direction of deviation of the growth laws from linearity during slow growth (when using equation (2)). This deviation is shown in the heatmaps using the ratio of the slope of growth laws (Figure 5a) when the birth rate is low ($\epsilon_P$ is 0.1 higher than the value that results in a zero birth rate) over when the birth rate is high ($\epsilon_P = 20$). Different panels show different recycling efficiencies. If this ratio exceeds 1, the growth laws are concave down, and if the the ratio is between 0 and 1, the growth laws are concave up, similar to the observations. This figure shows that the ratio is always larger than one, meaning the growth laws are always concave down. Other parameter values: $\epsilon_B = 3$, and $\phi_S = 0.5$

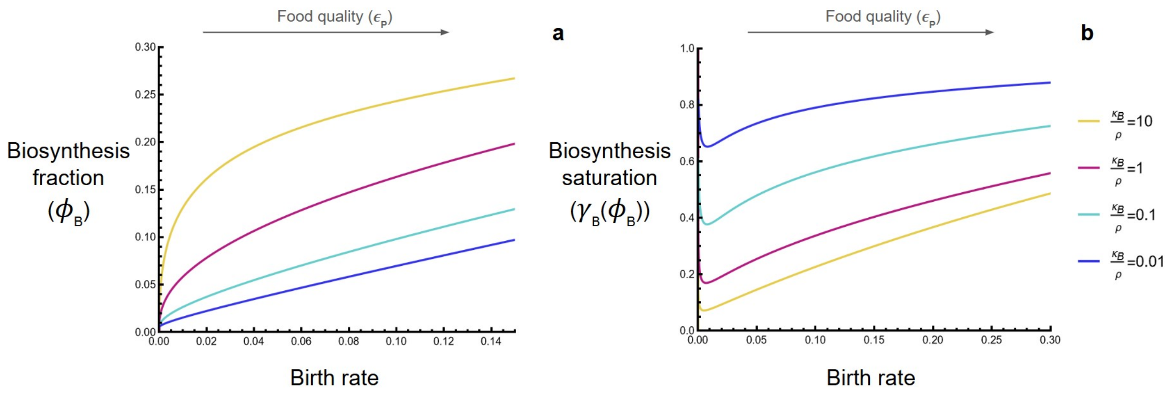


Figure S7: When using equation (S3), the predicted shape of bacterial growth laws becomes more linear when the macromolecular density is larger than the half-saturation constant of the biosynthesis fraction ($\rho > k_B$). Other parameter values: $\phi_S = 0.5$, $c = 0.8$, $\epsilon_B = 2$, $k_B = 1$, $S_M = 0.005$, $D = 0.01$, and $\rho = 10$
